# DeepPath: Overcoming data scarcity for protein transition pathway prediction using physics-based deep learning

**DOI:** 10.1101/2025.02.27.640693

**Authors:** Yui Tik Pang, Katie M. Kuo, Lixinhao Yang, James C. Gumbart

## Abstract

The structural dynamics of proteins play a crucial role in their function, yet most experimental and deep learning methods produce only static models. While molecular dynamics (MD) simulations provide atomistic insight into conformational transitions, they remain computationally prohibitive, particularly for large-scale motions. Here, we introduce DeepPath, a deep-learning-based framework that rapidly generates physically realistic transition pathways between known protein states. Unlike conventional supervised learning approaches, DeepPath employs active learning to iteratively refine its predictions, leveraging molecular mechanical force fields as an oracle to guide pathway generation. We validated DeepPath on three biologically relevant test cases: SHP2 activation, CdiB H1 secretion, and the BAM complex lateral gate opening. DeepPath accurately predicted the transition pathways for all test cases, reproducing key intermediate structures and transient interactions observed in previous studies. Notably, DeepPath also predicted an intermediate between the BAM inward- and outward-open states that closely aligns with an experimentally observed hybrid-barrel structure (TMscore = 0.91). Across all cases, DeepPath achieved accurate pathway predictions within hours, showcasing an efficient alternative to MD simulations for exploring protein conformational transitions.

## Introduction

Proteins are fundamental to a plethora of biological processes, with their functions intrinsically linked to their three-dimensional structures. Accurate predictions of these structures have long been a central challenge in computational biology. The advent of deep learning techniques has revolutionized this field, particularly with the development of AlphaFold2 (AF2),^1^ swiftly followed by tools such as RoseTTAFold,^2^ ESMFold,^3^ AlphaFold3^4^ and others.^5–8^ These methods enable the accurate prediction of static protein structures from amino acid sequences, providing researchers with invaluable tools to investigate protein structures, functions, and interactions at an atomistic level.^9–13^ However, proteins are not static entities; they are dynamic molecules that undergo conformational changes essential for their biological functions.

Protein dynamics are essential to enable multiple functional states, ligand interactions, and participation in complex cellular pathways.^14–17^ Insights into these conformational changes are vital not only for basic research, but also for practical applications like drug discovery. Understanding conformational flexibility can guide the rational design of therapeutic molecules for efficacy and selectivity.^18,19^ MD simulations are traditionally the gold standard for studying protein dynamics in silico. They offer atomistic insights into molecular movements based on thermodynamic principles.^20^ However, these simulations are computationally demanding. Even with state-of-the-art supercomputers, MD typically achieves only nanoseconds to microseconds per day,^21–23^ far short of the milliseconds-to-seconds timescales required for many biological processes. Enhanced sampling techniques^24–29^ and recent machine learning (ML) integrations^30–34^ aim to address this limitation, but they often remain computationally expensive and require highly specific input settings for different systems.

Building on the success of ML methods in static protein structure prediction, researchers have explored ways to leverage these tools to generate more diverse conformations.^35–39^ For example, masking coevolutionary signals in multiple sequence alignment (MSA) inputs^36–38^ or introducing alanine mutations^35,40^ were shown to induce some degree of conformational diversity in the generated structures.^40^ Other strategies include retraining models with additional structures derived from long-timescale MD simulations,^41^ adding diffusion-based or flow-matching-based auxiliary networks,^39,42,43^ or designing tailored architectures and training data for a smaller subset of proteins.^44–46^ These methods have achieved varying degrees of success. However, a persistent challenge shared by all is the limited availability of training data for dynamical protein structures, which potentially limits their ability to generalize effectively.^42^

Active learning (AL), an ML strategy designed to overcome data scarcity, offers a promising yet under-explored solution to this problem. Traditional supervised learning relies on large and static datasets. On the other hand, AL focuses on identifying data points where the model is uncertain and labeling them by interacting with an oracle (e.g., a human expert or physics-based computational model).^47^ This approach has been successfully applied in fields such as medical image diagnosis,^48–50^ autonomous driving,^51–54^ and molecular chemistry,^55–58^ drastically reducing the amount of labeled training data required. In the context of protein dynamics, AL holds the potential to address data scarcity by providing a reliable framework to train models specifically for proteins of interest, especially when paired with a robust oracle capable of evaluating structural reliability across diverse conformational landscapes.

In this paper, we introduce DeepPath, a novel AL algorithm for generating physically reliable protein transition pathways between two given structures. DeepPath leverages a molecular mechanics (MM) energy evaluator as an oracle to iteratively generate and validate intermediate structures without prior knowledge of the transition. To evaluate its capabilities, we tested DeepPath on a number of biologically relevant protein systems that exemplify key challenges in protein dynamics and large conformational changes. These systems range from soluble to membrane proteins, and from non-linear domain motion to substrate secretion. Specifically, we examined three protein transitions: activation of Src homology 2-containing protein tyrosine phosphatase-2 (SHP2), contact-dependent growth inhibition B (CdiB) helix 1 expulsion, and opening of the β-barrel assembly machinery (BAM) complex. We show that DeepPath efficiently explored the conformational spaces across all test cases, generating low-energy transition pathways within hours of training. Its predictions not only match existing data but also reveal novel conformations, broadening our understanding of these systems. Overall, these results underscore the potential of AL as a scalable approach for predicting dynamic protein structures with all-atom resolution, even in the absence of large training datasets.

## Results

### Model overview

DeepPath incorporates an AL framework to autonomously generate training data through interacting with a molecular mechanical (MM) energy minimizer (Fig. 1a). This approach directly addresses the scarcity of intermediate structures in public protein databases by self-generating training data on a per-protein basis, instead of relying on static training datasets. Unlike pool-based AL where the model selects data out of some databases for labeling, DeepPath generates novel structures and validates them using an MM energy minimizer as an “oracle.” To initiate the pathway exploration, short MD simulations are conducted of each of the input end-states. These initial training data need not cover any of the transition states, but rather they give clues to the network about the relative mobility of each part of the protein. Then, DeepPath starts generating its own training data iteratively by constructing potential intermediate structures and validating their energies via a MM energy minimizer, slowly broadening the range of generated conformations while ensuring these structures remain realistic. Through this iterative process, DeepPath searches different possible transition pathways, expanding its training database until no alternative pathways with lower energies can be found.

**Figure 1:**
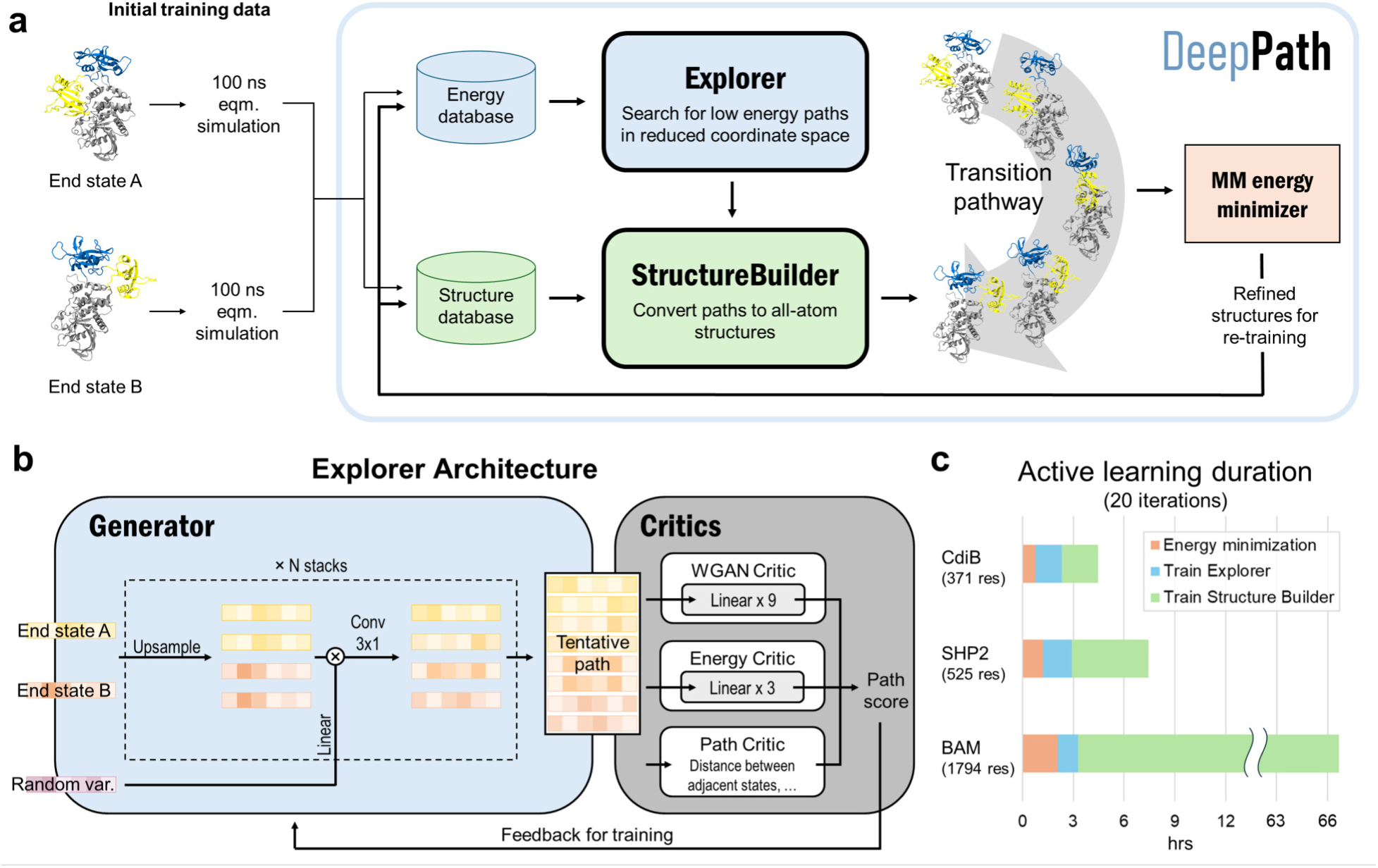
DeepPath Overview. a, AL routine for training DeepPath. First, short MD equilibrium simulations are performed at each of the end state to generate some initial training data. This data is utilized to train the two modules of DeepPath, the Explorer and Structure Builder, which together produce all-atom structures along tentative transition pathways. Selected generated structures are refined with their corresponding energies determined using the MM energy minimizer. These refined structures are added to the training database to re-train DeepPath and gradually improve the reliability of the transition pathways generated as the training cycle continues. b, The Explorer is a modified progressive GAN, with a combination of critics that work together to ensure the paths generated are realistic and energetically feasible. c, Training duration for 20 iterations of AL using one GPU for three protein test cases: CdiB, SHP2, and the BAM complex.

The architecture of DeepPath comprises two specialized neural network modules: the Explorer and the Structure Builder (Fig. 1a). The Explorer is a progressive generative adversarial network (GAN)^59,60^ designed to predict an ensemble of transition pathways represented in a reduced coordinate set. The Structure Builder, on the other hand, is a custom neural network that translates the Explorer’s reduced coordinate outputs into detailed all-atom structures. The choice of the reduced coordinate set is versatile, provided it can effectively differentiate between distinct protein conformations. To ensure the model’s generality, we have selected all residue-residue pairwise distances that show significant variation beyond a predetermined threshold as our reduced coordinates for this implementation.

The modular design of DeepPath is crucial to its success. The use of a reduced coordinate set isolates the Explorer from the high-dimensional coordinate space, which is characterized by a complex, uneven energy landscape that complicates the exploration and optimization of conformations. Additionally, this modular approach simplifies the overall network architecture, reduces the volume of training data required, and enables rapid production of valid transition pathways.

From our testing, DeepPath generates reliable transition pathways within 20 AL iterations, which takes only a few hours on a single GPU for an average-sized protein (Fig. 1c) and achieves approximately 80% of the energy reduction observed at full convergence. For example, DeepPath computed the expulsion process of the H1 helix from the CdiB β-barrel, which includes 371 residues, in less than 4.5 hours. DeepPath also proved scalable by accurately predicting the transition pathway between the inward-and outward-open states of the 1794-residue BAM complex, albeit requiring a longer computational time of 66.7 hours. A breakdown of the training time reveals that the majority of training cost increment is allocated to the training of the Structure Builder, the largest network within DeepPath. The use of the frame aligned point error (FAPE) in its loss function contributes to a computational complexity that scales quickly with the size of the protein, increasing from merely 2.1 hours for CdiB to 63.4 hours for the BAM complex. The time spent on energy minimization also increases with the complexity and size of the protein. In contrast, the training time for the Explorer remains largely constant across different protein sizes, highlighting the efficiency achieved through the use of reduced coordinate sets.

### Explorer

The core innovation of DeepPath resides in the Explorer, a GAN responsible of exploring the vast protein conformational space and constructing plausible transition pathways between the two specified input end states. These pathways are represented as a series of pairwise distance arrays (x) that encode each intermediate state along the pathway. We chose progressive GAN as the fundamental architecture for the Explorer as we found its incremental nature to be highly compatible with our AL protocol (Fig. 1b). Progressive GAN grows the network progressively by starting the training at a low resolution, then adds new layers that model increasingly fine details as training progresses. We adopted this design for the Explorer by first training it to generate transition pathways represented by only four intermediate states, and then adding new layers to the generator and doubling the path resolution every five AL training iterations until a sufficient resolution is reached. This staged learning approach allows the network to first capture a broad overview of the energy landscape before fine-tuning the pathways to minimize energy more effectively, balancing global exploration with local optimization.

Usually, GANs are composed of a generator and a critic, with the generator trained to generate outputs that are deemed realistic by the critic, and the critic trained to distinguish the generated outputs from the real ones. These two networks are trained against each other to improve each other’s performance. However, requiring a priori pathway examples as training data defeats the purpose of DeepPath. Instead, the Explorer’s generator is trained against a collection of critics.^61^ Two of them are neural-network-based (*C*_WGAN_ and *C*_energy_), which judge the fitness of each individual intermediate states that makes up the pathways. The other one is math-based (*C*_path_), which evaluates if the intermediate states collectively form a smooth and continuous pathway. Hence, the loss function for the generator is:

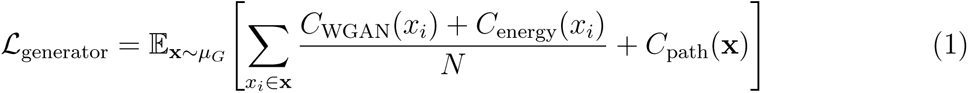

where N is the number of intermediate structures sampled from each generated path. *C*_WGAN_, implemented based on Wasserstein GAN (WGAN) with gradient penalty,^62^ is trained to distinguish between the pairwise-distance arrays produced by the generator against those from the structure database, guiding the generator to produce pairwise-distance arrays that resemble ones of the validated intermediate states. *C*_energy_ is trained to predict a structure’s energy from its pairwise-distance representation, serving to guide the generator towards low-energy intermediate states. While *C*_WGAN_ is trained together with the generator, *C*_energy_ is trained independently at each AL loop against the energy database. The three critics together ensure the generated pathways are realistic, low-energy, and continuous.

Outside of providing guidance for the generator, the *C*_WGAN_ and *C*_energy_ also serve another important purpose in the AL process. Due to the high computational cost of performing energy minimization (compared to the rest of the training loop), only a fraction of generated intermediate structures were energy minimized and had their energies evaluated. Drawing parallels to the AL criterion “query-by-committee”,^63^ we pick structures that obtain conflicting verdicts from *C*_WGAN_ and *C*_energy_. Specifically, if *C*_WGAN_ predicts the structure to be unrealistic but *C*_energy_ predicts it to be low energy, the structure is sent to have its energy evaluated. Conceptually, this criteria encourages structures that are rated novel and low-energy to be evaluated and added to the database, iteratively improving the training data quality and eventually leading to the generation of the most plausible transition pathways.

### Test case 1: SHP2

To evaluate DeepPath in a real-world scenario, we applied it to a protein target characterized by a significant domain shift that is crucial to its pathogenic behavior. Src homology-2 domain-containing protein tyrosine phosphatase-2 (SHP2) is a protein target that has been implicated in cancer but long considered undruggable.^64,65^ In a healthy cell, SHP2 is conventionally remains in its closed state and only transitions into its open state upon binding with its partner protein. In cancerous cells, SHP2 can go from the closed to open conformation spontaneously, without binding of the partner protein.^66,67^ In addition, the protein sometimes has point mutations that bias it to the open state.^68,69^ The overactive SHP2 in the open conformation results in cancerous cell proliferation, tumor invasion, and metastasis.^70,71^ While there have been numerous small molecules developed against SHP2, none are FDA-approved, and all are designed to target the closed state.^67,72^ By understanding the transition from the closed to the open state, more effective therapeutics can be designed to inhibit this conformational transition and, thus, the pathogenesis of cancer.^70^ These conformational changes are particularly interesting for assessing whether DeepPath can generate pathways that are not strictly linear but instead rotational, and whether it can prevent the N-SH2 domain from colliding with the PTP domain. We were particular interested in the E76A variant due to the partially open state that had been previously reported.^68^

DeepPath was initially trained on two 100-ns equilibrium MD trajectories, one originating from the inactive state and the other from the active state. Due to the short length of these trajectories, only a limited conformational space near the starting end-states was sampled. Following the initial training, DeepPath was trained for 50 iterations using the AL protocol. In each iteration, DeepPath evaluated the validity of selected structures it generated by determining whether their MM potential energy, computed by the energy minimizer, fell below an adaptive threshold. Once 32 valid structures were identified, they were incorporated into a database as part of the training dataset. Over the whole course of the AL process, a total of 1600 new structures were added to the training set.

Fig. 2a shows the SHP2 transition pathway predicted by DeepPath. Consistent with previous studies,^66,67^ DeepPath identified a pivotal movement of the N-SH2 domain, which underwent a hinge-like swing around the PTP domain during the transition between the active and inactive states. Simultaneously, DeepPath predicted a rotation of the C-SH2 domain, positioned on top of the PTP domain. Interestingly, despite the large overall rootmean-square-deviation (RMSD) between the active and inactive states, the RMSD of each individual domain remains low throughout the predicted pathway. This suggests that Deep-

**Figure 2:**
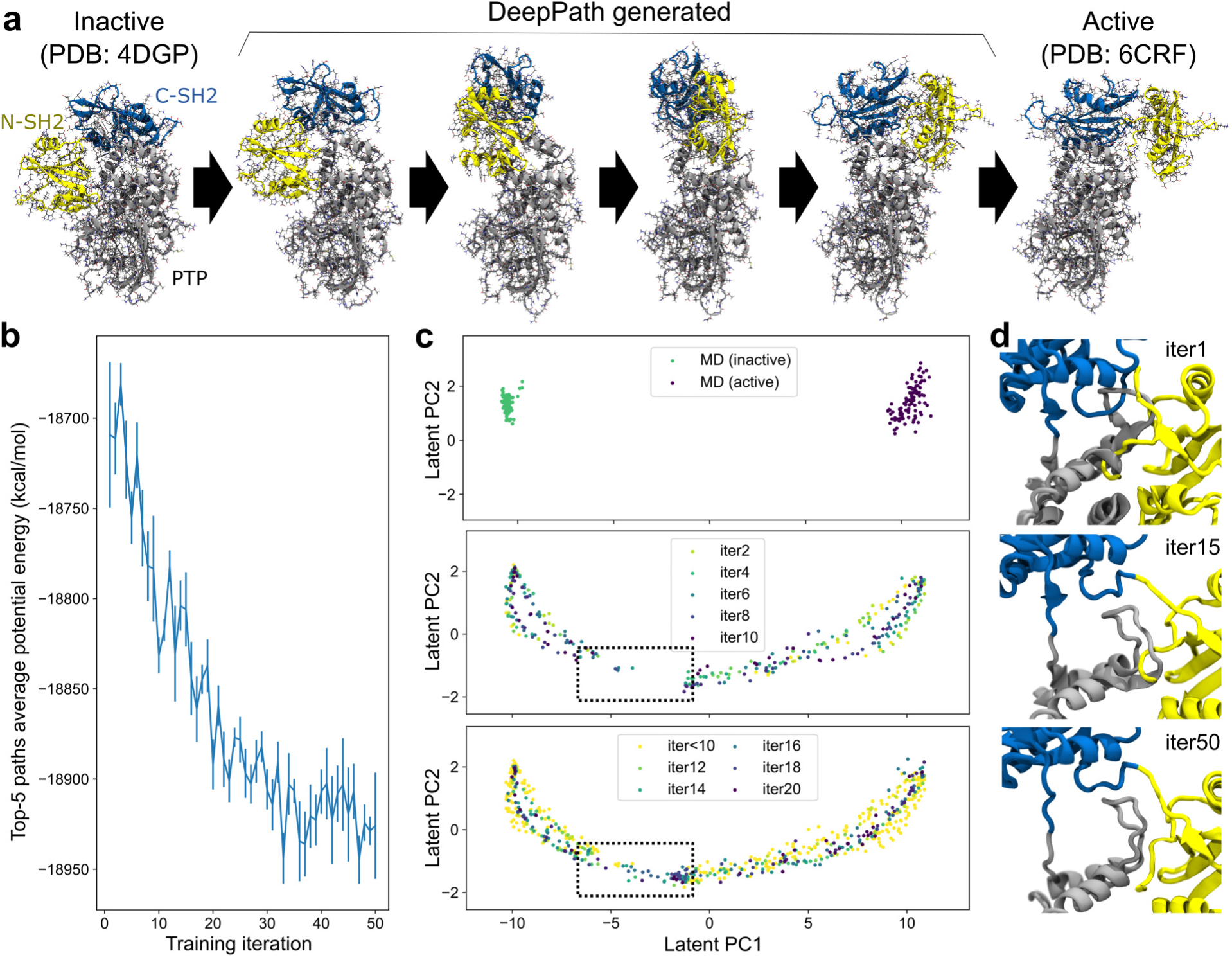
DeepPath predicted the transition paths between SHP2 active and inactive states. a, The predicted transition pathway of SHP2 highlights the rotational movement of the NSH2 domain around the PTP domain, accompanied by a rotation of the C-SH2 domain. b, The average potential energy of the top-5 generated pathways decreases as training proceeds. c, Training structures generated at different stages, projected onto the principal components of their reduced representations. Initial equilibrium simulations are shown in the top panel, AL iterations 1–10 are shown in the middle panel, and iterations 11–20 in the bottom panel. The dotted square highlights the gap in the pathway that occurred early in training, which was successfully filled after 15 iterations of AL training. d, Close-up views highlights the challenge of generating structures in the region where the pathway initially breaks. Top: Before training, DeepPath predicts a structure with steric clashes around the A236–K244 loop from the PTP domain. Middle: After 15 iterations of AL, DeepPath lifts the C-SH2 domain, creating space for the A236–K244 loop to maneuver under the linker between the C-SH2 and N-SH2 domains, resolving the major steric clash between the A236–K244 loop and the C-SH2 domain. Bottom: At convergence, DeepPath completely resolves the steric clash issues, preventing the C-terminus from hooking into the A236–K244 loop.

Path preserved the local structural integrity of SHP2 during the transition, avoiding distortions that could compromise its functional state. Moreover, along the predicted transition pathway, DeepPath identified an intermediate conformation that closely resembles the partially open state reported by Tao et al., notably recovering a critical contact between residues Y63 and E508.

Examining the progression of DeepPath during the AL protocol, we observed that the average potential energy of the top-5 generated pathways decreases gradually during the training and stabilizes at approximately –18,920 kcal/mol (Fig. 2b). This drop in energy can be explained by analyzing the structures generated by DeepPath during the training process. Early in the training, DeepPath showed an “understanding” that the N-SH2 domain must rotate around the PTP domain to transition between the active and inactive states. However, it initially lacked the ability to determine the optimal distance between the NSH2 and PTP domains, resulting in steric clashes and higher energy barriers. Specifically, a loop (residues A236–K244) from the PTP domain clashed with both the N-SH2 and CSH2 domains (Fig. 2d). When examining the validated structures in the training database, we noticed an early gap in the pathway, where all of the generated structures within the gap failed to obtain potential energies below the required threshold (Fig. 2c). However, DeepPath successfully generated valid structures that bridged this gap by the 14th iteration. It continued to produce more low-energy structures in subsequent iterations, eventually filling the gap entirely. The final predicted pathway reveals that DeepPath adjusted the C-SH2 domain, lifting it to create more space for the A236–K244 loop to pass underneath the linker between the C-SH2 and N-SH2 domains (Fig. 2d).

### Test case 2: CdiB

DeepPath’s application extends beyond proteins with multiple stable conformations. In this section, we demonstrate its capability to study the expulsion of a domain from a transmembrane channel. The contact-dependent growth inhibition (CDI) system is a mechanism utilized by Gram-negative bacteria to suppress the growth of neighboring bacterial cells to reduce competition.^73^ As a member of the Two-Partner Secretion (TPS) family, the CDI system consists of two components: the CdiA toxin and the CdiB transporter. Here, we focused on CdiB, a β-barrel protein that inserts into the bacterial outer membrane, allowing CdiA to pass through its lumen and be displayed on the cell surface.^74–77^ However, in its resting state, this β-barrel is occluded by an N-terminal α-helix (H1) during activation, H1 exits first into the periplasm to enable CdiA secretion.^74,78–80^

We used DeepPath to generate the exit pathway of the H1 helix and compared our results with a previous study^80^ that explored this process using steered MD (SMD) and replica exchange umbrella sampling (REUS). In the resting state, the upper half of H1 was tightly packed within the barrel lumen’s narrower, more constricted region, limiting its movement. As expulsion progresses, H1 gradually slides downward, freeing itself from these constraints, and makes transient interactions with residues in the lower section of the β-barrel (Fig. 3b). Further examination of the trajectories revealed a particularly prominent interaction between H1 and the β4–β5 region.

**Figure 3:**
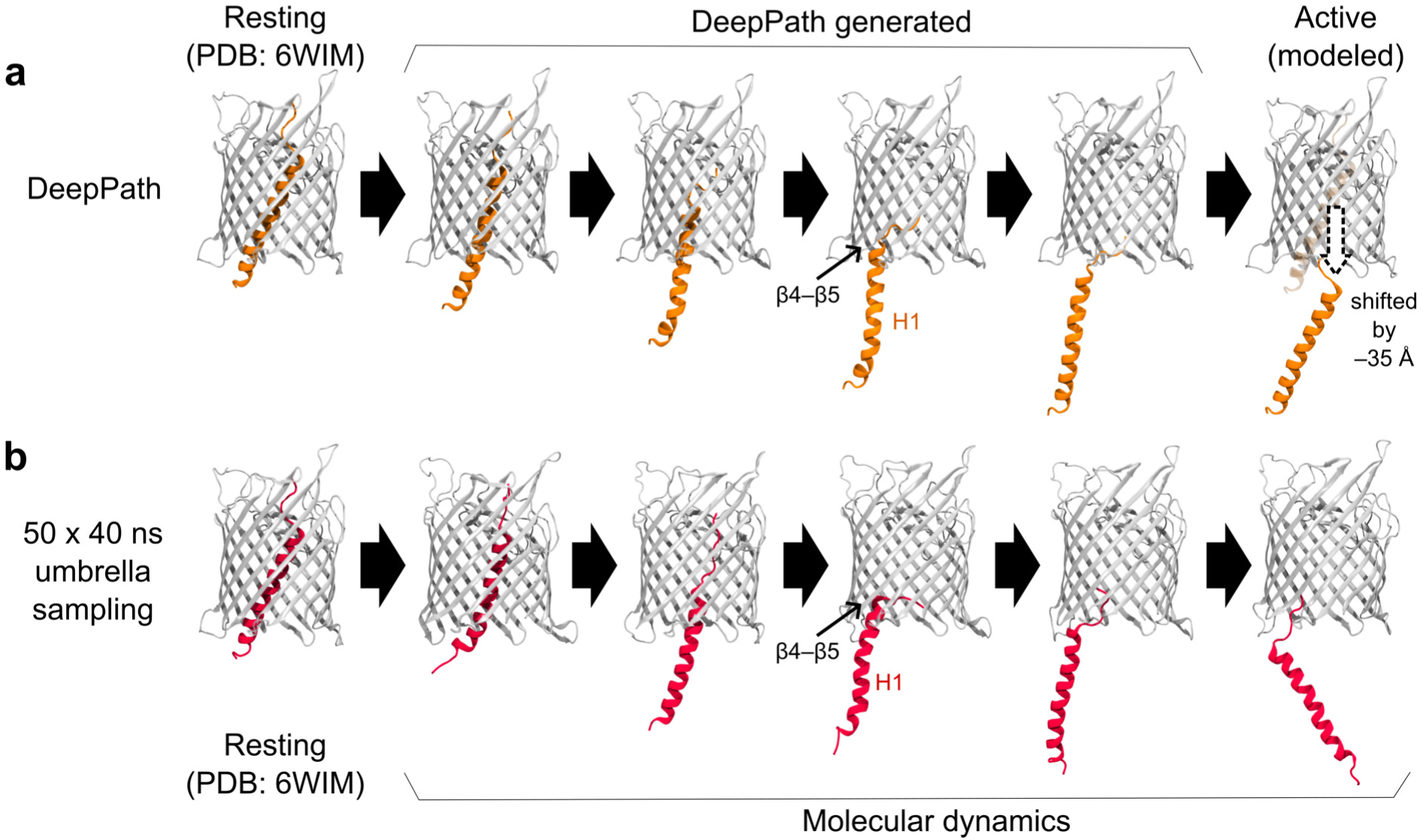
Exit pathway of the N-terminal α-helix (H1) in CdiB as predicted by a, DeepPath and b, REUS simulations from a previous study.^80^ DeepPath efficiently generated exit pathways that closely match the REUS-derived pathway, despite requiring significantly less computational time. Notably, DeepPath accurately captured the key transient interaction between H1 and β4–β5 during expulsion.

Here, we assessed whether we can use DeepPath to recover this pathway with only the resting structure known. To construct the active state, we shifted H1 downward by 35 Å using VMD, positioning it entirely within the periplasmic space. Two 10-ns equilibrium MD simulations were run, one starting from the resting state and one from the modeled active state. Additionally, to aid DeepPath in generating an initial guess, a series of modeled intermediate structures were generated by incrementally displacing H1 downward at an interval of 0.35 Å. These additional structures underwent energy minimization but were not subjected to further simulations.

The H1 exit pathway predicted by DeepPath after 50 iterations of AL training highly resembles the one identified by the 2-µs aggregated REUS simulations (Fig. 3). The model predicted that H1 initially straightened itself slightly as it exits the narrow upper section of the β-barrel before moving downward, where it established transient interactions with the periplasmic end of β4–β5. Finally, it detached H1 from the β-barrel, reaching the modeled active conformation. Over the course of training, the interaction energy between H1 and the β-barrel gradually decreased (Fig. 4a). This trend suggests that DeepPath systematically explored alternative orientations of H1 and identified those leading to more favorable interactions.

**Figure 4:**
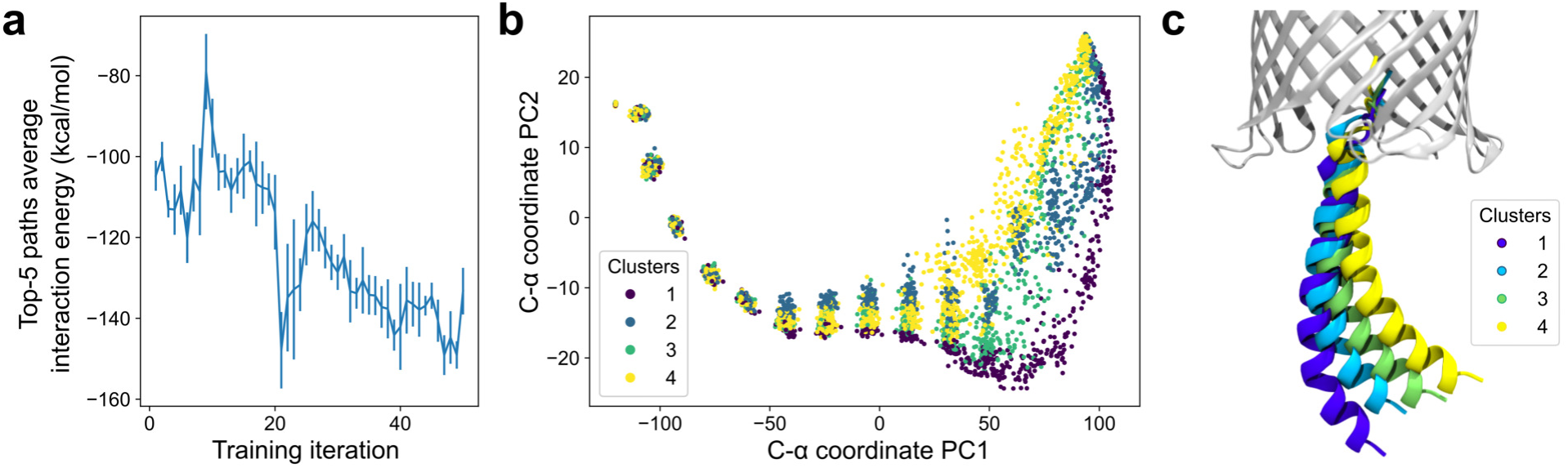
Diversity and energy refinement of DeepPath-generated H1 exit pathways. a, Evolution of the interaction energy between H1 and the β-barrel over AL training iterations shows an overall reduction in energy as DeepPath refined its predicted pathways. b, PCA of the generated pathways based on the C-α coordinates of H1, with the β-barrel aligned. Each point represents an intermediate conformation, colored according to pathway clusters identified via path similarity analysis (PSA). c, Representative conformations from each pathway cluster, illustrating structural variability in the periplasmic region while maintaining key contacts with the β-barrel.

Beyond reproducing the H1 exit pathway, DeepPath demonstrated the ability to generate diverse pathway predictions, particularly in regions where structural diversity does not come with an energy penalty. As observed in the REUS simulations, once H1 exited the β-barrel, its unconstrained portion could adopt a wide range of conformations. DeepPath also captured this change of flexibility. Looking at an ensemble of 256 transition pathways generated after training, all pathways were highly consistent when H1 was still confined within the barrel. However, as more residues emerged into the periplasmic space, the diversity among the generated pathways greatly increased while still maintaining key interactions, such as those between H1 and β4–β5 (Fig. 4b,c).

To further assess the extent of this diversity, we applied path similarity analysis (PSA)^81^ to cluster the pathway ensemble generated by DeepPath. We identified four distinct pathway clusters using a cluster distance cutoff of 28 in Ward criteria, indicating multiple viable exit pathways predicted by DeepPath. These clustering results align with the principal component analysis (PCA) of the H1 C-α coordinates, where the four pathway groups well-separated along the first two principal components (Fig. 4b). Notably, despite the structural variation, the average potential energies among all pathway clusters remained consistent at ∼12,075 kcal/mol, confirming that the pathway diversity was achieved without sacrificing energy efficiency. These findings suggest that DeepPath’s prediction extends beyond a single transition pathway and instead approximates a transition tube, encompassing a diverse set of thermodynamically accessible routes at finite temperature.

### Test case 3: the BAM complex

To assess the scalability of DeepPath, we applied it to the BAM complex from *Escherichia coli*, a large multi-protein system with nearly 2,000 total residues. The BAM complex consists of the transmembrane BamA β-barrel and the accessory proteins BamB-E. It plays a crucial role in the biogenesis of outer-membrane proteins (OMPs) in Gram-negative bacteria by facilitating their folding and insertion into the outer membrane.^82–84^ BAM adopts two distinct conformations: the inward-open and outward-open states.^85–89^ In the inward-open state, the lumen of BamA’s β-barrel is accessible from the periplasm, allowing nascent OMPs to enter. In contrast, the outward-open state closes this passage while separating the first (β1) and last (β16) strands of the β-barrel, forming the lateral gate (LG) – a critical entry point for nascent OMPs to integrate into the membrane. Recent structural studies revealed that during OMP insertion, BAM transiently forms a hybrid-barrel intermediate, where the substrate OMP uses BamA’s β1 strand as a scaffold for folding.^90–94^ Here, we used DeepPath to predict the transition pathways between the inward- and outward-open states absent a substrate, training it solely on structural data from two 500-ns equilibrium MD simulations initiated from these two endpoint structures. Strikingly, DeepPath’s predicted transition pathway revealed an intermediate conformation in agreement with an experimentally resolved hybrid-barrel structure of BAM interacting with EspP, a substrate OMP.^90^

From the DeepPath-predicted transition pathway, it is clear that conformational changes in BamA’s β-barrel are tightly coupled with a counterclockwise rotation of the periplasmic domains (Fig. 5a). Between the inward- and outward-open states, DeepPath predicted a two-stage transition in the β-barrel: (1) LG opening and (2) base narrowing. During the first half of the transition, the β1 strand moved in sync with the rotation of the periplasmic domains. This motion quickly broke the backbone hydrogen bonds between β1 and β16 in the inward-open state, initiating LG opening. As β1 continued to straighten and peel away from β16, the LG reached its maximum opening width after the periplasmic domains rotated by approximately 15° (Fig. 5b). In the second half, the periplasmic domain rotation correlated with the narrowing of the β-barrel’s base. The transition was complete once the periplasmic domains rotated by approximately 40° in total, at which point the base of the β-barrel constricted, fully sealing the periplasmic opening and the outward-open state was stabilized.

**Figure 5:**
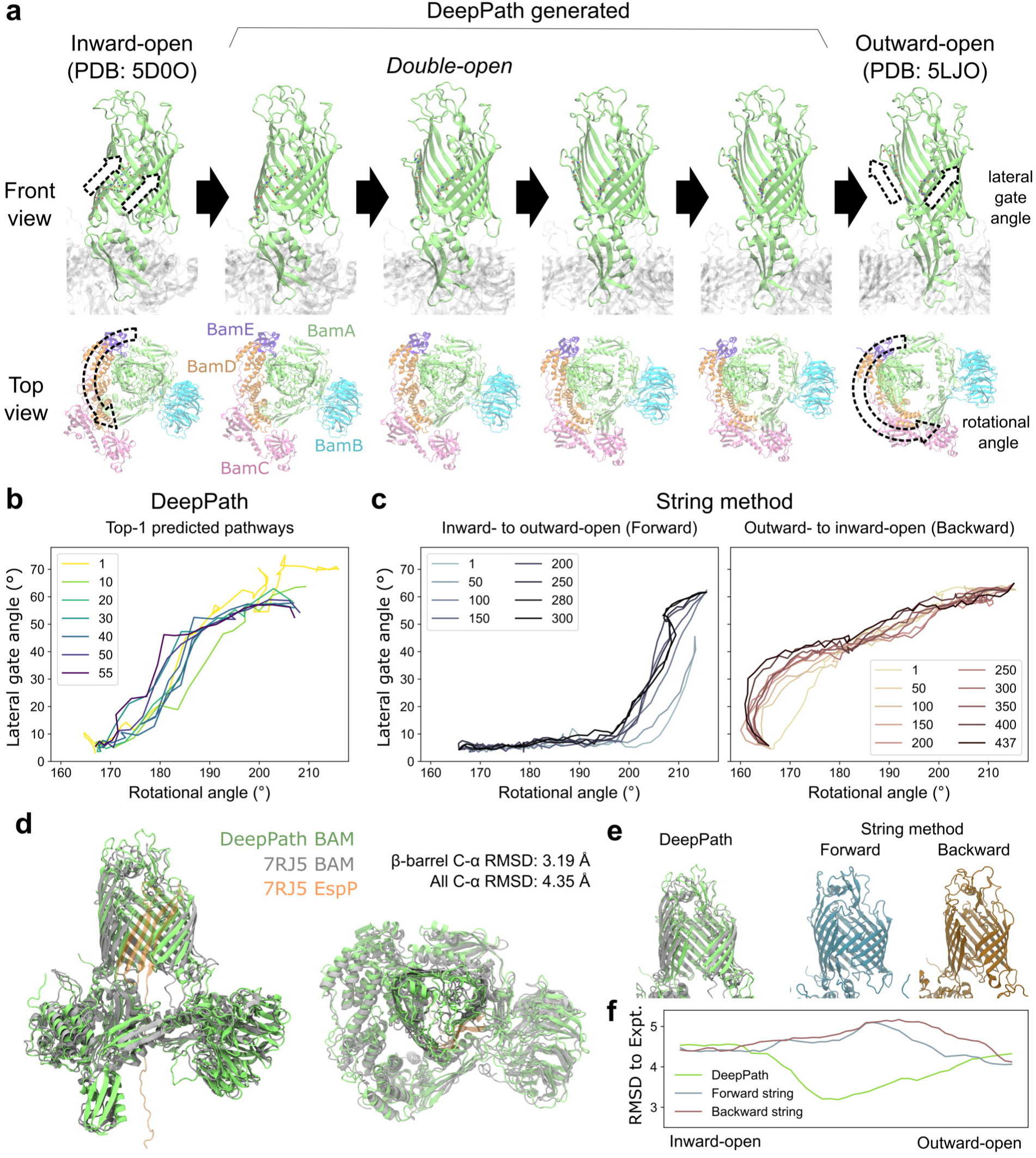
DeepPath-generated transition pathway of the BAM complex from the inward-to outward-open state and comparison to the string method. a, Snapshots from the final iteration of DeepPath results showcasing the angular opening of the lateral gate (LG), the movement of the P5 domain from left to right, and the counter-clockwise rotation of the periplasmic domains BamB–E as the LG opened. b-c, Evolution of the transition pathways as represented by the LG angle and periplasmic domain rotational angle. b, Top-1 pathways predicted by DeepPath. c, Pathways obtained using SMwST, depicted for both inward-tooutward-open (forward) and outward-to-inward-open (backward) transitions. d, The DeepPath-predicted “double-open” structure (green) overlapped with the cryo-EM structure of BAM/EspP hybrid-barrel intermediate (gray for BAM and orange for EspP; PDB: 7RJ5). Only shared residues of BAM were shown. e, Comparison of the β-barrel structure predicted by DeepPath and SMwST during transition. f, C-α RMSD between the predicted β-barrels and the hybrid-barrel experimental structure along the predicted transition pathways by DeepPath and SMwST.

The DeepPath prediction suggests that BAM adopts an intermediate “double-open” state, where the BamA β-barrel lumen is accessible from both the periplasmic space and the membrane simultaneously. This observation is reminiscent of the hybrid-barrel intermediate, where both the LG and the periplasmic entrance must open to allow the nascent substrate OMP to pass through the BamA lumen and reach the LG as folding proceeds. To test whether the DeepPath-generated double-open structure corresponds to such an intermediate, we compared it to an experimentally resolved BAM/EspP hybrid-barrel intermediate structure (PDB: 7RJ5).^90^ Despite the absence of the substrate OMP in DeepPath’s model, the alignment of the BamA β-barrel is remarkably high (Fig. 5d). The C-α RMSD between the predicted and experimental structures measured only 3.19 Å and a TM-score of 0.86. This structural agreement also extended to the membrane-inserting hydrophobic loops. Even without explicitly modeled membrane lipids, DeepPath correctly oriented the membrane-inserting hydrophobic loops, proving its capability to infer and preserve structural constraints purely from MD training data. Across all shared residues of the BAM complex, the C-α RMSD was 4.35 Å with a TM-score of 0.91. This demonstrates that DeepPath accurately captured the global structure, including the relative domain arrangement of the accessory proteins BamB-E (Fig. 5d).

As a comparison, we also applied the string method with swarms of trajectories (SMwST)^95^ to compute the same inward-to-outward-open transition pathway of the BAM complex. To generate the initial pathway estimates, we first ran two separate targeted MD (TMD) simulations: one pulling BAM from the inward- to outward-open state (forward) and another pulling it from the outward- to inward-open state (backward). These TMD-derived pathways were then used to seed two independent SMwST calculations. Each string was discretized into 51 equally spaced images, and 440–640 ps simulations were run per image, per iteration to evaluate the local energy gradients and update the pathway. The forward and backward strings were optimized for 300 and 437 iterations, respectively, accumulating to a total simulation time of 14.5 µs.

Overall, SMwST made significant progress in smoothing out abrupt transitions and reducing fluctuations, but major structural defects inherited from the original TMD pathways remained. Specifically, the forward TMD struggled to break the existing interactions that lock the LG in the inward-open state, while the backward TMD delayed widening the base of the β-barrel until the final stages of the pathway. Even after hundreds of SMwST iterations, remnants of these defects persisted in the optimized pathways. Notably, the forward and backward strings follow markedly different routes. In the forward string, the LG remains closed until the periplasmic domain has rotated for more than 30°, whereas in the backward string, LG closure occurred before any periplasmic domain rotation (Fig. 5c). Additionally, the forward transition retained a pronounced kink in the β1–β2 region, a distortion carried over from the initial TMD trajectory (Fig. 5e). Most importantly, neither the forward nor backward SMwST-generated pathways predicted the “double-open” state that DeepPath did. In fact, all intermediate structures generated by SMwST exhibited higher β-barrel C-α RMSD against the experimental hybrid-barrel structure (PDB: 7RJ5) than the starting end states, whereas DeepPath’s predictions consistently achieved lower RMSD than the starting structures (Fig. 5f). This highlights DeepPath’s superior predictive power in uncovering unseen intermediate structures accurately.

Comparing the optimization process of DeepPath with that of SMwST, we found that DeepPath converged much faster, both in terms of iterations required and computational cost. A key advantage of DeepPath is that, unlike traditional simulation-based methods, it is not constrained by directionality. Regardless of the simulation methods used to seed SMwST, the process must begin from one state and apply biasing forces to drive the system toward the other state. This approach unavoidably introduces distortions and human bias into the generated pathway. Once introduced, it would require hundreds of nanoseconds or even microseconds of equilibrium simulations to relax the structures and eliminate the defects, making it challenging for SMwST to recover physically realistic intermediates. In contrast, DeepPath generates all intermediate conformations independently and simultaneously, without relying on biasing forces. Instead, it explores conformational space using a neural-network-driven approach that leverages the protein’s intrinsic degrees of freedom and iterative energy evaluations, producing physically plausible initial guesses that require minimal structural corrections. Beyond better initial structure generation, DeepPath also demonstrated a strong ability to repair secondary structure distortions over optimization iterations. In early iterations, we observed twisting and breaking of native hydrogen bonds between BamA’s β1 and β2, as well as disruptions in the secondary structure of extracellular loop 1. However, as the active learning iterations progressed, DeepPath gradually corrected these defects, ultimately producing secondary structures that align with the stable input conformations.

## Discussion

In this manuscript, we introduce DeepPath, a novel deep learning approach that generates low-energy transition pathways using AL, eliminating the need for large pre-existing training datasets. Instead, DeepPath predicts transition pathways by iteratively generating candidate conformations and assessing their physical feasibility using all-atom MM force fields. Through repeated structural refinement and exploration based on this energy evalu ation, DeepPath gradually builds up the training dataset and efficiently produces atomistic-resolution transition pathways within hours.

DeepPath demonstrates how AL can be leveraged to exploit the physical and chemical knowledge embedded in MM force fields, training deep learning models to generate novel protein conformations beyond the limitation of data availability. Unlike a previous approach that directly incorporates force field energy terms into the loss function,^96^ our approach introduces an intermediate network, the Energy Critic, which learns an approximate energy landscape to guide DeepPath’s generation. This indirect learning strategy prevents instability caused by direct optimization on rugged MM force fields and significantly reduces computational cost by limiting the number of explicit energy evaluations. However, the current implementation only evaluates protein potential energies and accounts for environmental factors using implicit solvent rather than explicitly modeling water molecules or membrane lipids. Future work could address this limitation by extending the Structure Builder module to construct other biomolecules including water, lipids, and nucleic acids, enabling more accurate computation of interaction energies with the environment.

We evaluated DeepPath across three diverse cases of protein transitions: SHP2 activation, CdiB H1 expulsion, and BAM complex LG opening. Across all test cases, DeepPath successfully generated transition pathways that closely match previous studies, demonstrating its ability to produce accurate results through refinement via AL. Remarkably, DeepPath predicted a “double-open” intermediate conformation of the BAM complex, which closely aligns with the BAM/EspP hybrid-barrel structure resolved by cryo-EM,^90^ achieving a TMscore of 0.91. This was especially significant given the absence of the substrate OMP in DeepPath’s input and output. Beyond accuracy, DeepPath also demonstrated the ability to generate diverse transition pathways. In the CdiB test case, pathway clustering via PSA and PCA revealed four distinct transition route families and highlighted DeepPath’s capacity to explore multiple low-energy transition pathways. Additionally, DeepPath proved to be computationally efficient, identifying low-energy transition pathways within 4.5 hours for CdiB (371 residues) and 66 hours for the BAM complex (1794 residues) on a single GPU. These results highlight DeepPath’s ability to rapidly generate physically plausible transition pathways, making it a valuable tool for accelerating the exploration of conformational space. By efficiently identifying viable transition routes, DeepPath complements MD simulations, reducing the need for costly brute-force sampling that could take months – even with significantly greater computational resources. However, DeepPath’s predictive capability has some inherent limitations. In the CdiB H1 test case, while DeepPath accurately captured the strong electrostatic interactions between H1 and the β-barrel, it missed the hydrophobic interactions that was previously revealed by the extended (10-µs) REUS simulations. This is likely stems from DeepPath’s lack of explicit entropy consideration, making it less effective at predicting hydrophobic interactions and, in other cases, properly disfavoring extended structures with high entropic penalties.

It is important to note that DeepPath is designed to complement, not replace, MD simulations and enhanced sampling techniques, which remain the gold standard for capturing thermodynamic and kinetic properties. By rapidly generating plausible protein transition pathways, DeepPath can help guide MD sampling toward relevant regions and accelerate convergence in free energy calculations. Beyond its role in accelerating simulations, DeepPath also shows promise for drug discovery applications. Understanding conformational dynamics is critical for designing drugs that target allosteric sites or stabilize specific functional states of proteins. For example, many kinase inhibitors and molecular chaperone modulators rely on exploiting transient conformational states.^97–100^ DeepPath helps to identify these states, aiding in the development of more selective and effective therapeutics.

Beyond transition pathways, the AL framework presented here could be extended to a variety of biomolecular structure prediction challenges, including ligand-induced conformational changes, protein folding pathways, and RNA structure prediction. These problems face an even greater scarcity of high-quality structural data, making AL particularly valuable for efficiently navigating large conformational spaces and generating low-energy structures.

The AL framework introduced in this work effectively integrates ML with MM force fields, enabling structural prediction beyond the scope of the available training dataset. Future extensions of this approach could incorporate diffusion models or flow-based generative architectures, improving the ability to generate diverse structural ensembles, and better capturing the full conformational landscape of biomolecules beyond the current state-of-the-art supervised models.

## Acknowledgement

This project was partially supported by a Texas Advanced Computing Center (TACC) Frontera Fellowship. Frontera is supported by NSF grant OAC-1818253. This work was also supported by the National Institutes of Health (R01-GM148586). Additional computational resources were provided through ACCESS (grant TG-MCB130173), which is supported by National Science Foundation grants 2138259, 2138286, 2138307, 2137603, and 2138296. This work also used the Hive cluster, which is supported by the NSF under grant number 1828187 and is managed by the Partnership for an Advanced Computing Environment (PACE) at the Georgia Institute of Technology.

